# Unprecedented account of mortality and morbidity in free ranging Formosan Pangolin (*Manis pentadactyla pentadactyla*)

**DOI:** 10.1101/325167

**Authors:** Nick Ching-Min Sun, Bharti Arora, Jing-Shiun Lin, Wen-Chi Lin, Meng-Jou Chi, Chen-Chih Chen, Kurtis Jai-Chyi Pei

**Author notes:** These authors contributed equally to this work (Nick Ching-Min Sun, Bharti Arora). Correspondence: Kurtis Jai-Chyi, Pei, Institute of Wildlife Conservation, National Pingtung University of Science and Technology, Pingtung, Taiwan. 1, Shuefu Road, Neipu, 91201 Pingtung, Taiwan. #6590; Mail (N.C.M. Sun), (B. Arora), (J.S. Lin), (W.C. Lin), (M.J. Chi), (C.C. Chen). Present contact: Department of Veterinary Medicine, College of Veterinary Medicine, National Pingtung University of Science and Technology, Pingtung, Taiwan (Wen-Chi Lin).

## Abstract

Globally, pangolins are overt for poaching and illegal trade, but Taiwan projects totally a contrary image where their population is stable and increasing. This paper illustrated the factors responsible for causing mortality and morbidity in free ranging Formosan pangolin (*M. pentadactyla pentadactyla*). Results from radio-tracking showed that Formosan pangolins are highly susceptible to stuck in tree hallows or ground burrows despite being proficient burrowers, and killed by gin traps, especially during the dryer season. Whilst the data consolidated from the Pingtung Rescue Center for Endangered Wild Animals illustrated the trauma (73.0%) was the major reason of causing morbidity in Formosan pangolin. The gin traps were the leading cause of trauma (77.8%) along with tail injuries caused by dog attack (20.4%). Despite all the pressing data about the injuries Taiwan, it is able to establish substantial success rate in rescuing and releasing pangolins through consolidating and applying valuable information among the rescue centers in the span of two decades. Taiwan has made a phenomenal progress in sustaining a salubrious population of pangolin but the situation requires continuous examination to ensure the existence of this species on the island.

## Introduction

Formosan pangolin (*Manis pentadactyla pentadactyla*) is a subspecies of Chinese pangolin found only in the sub-tropical island Taiwan [1]. It occupies low to mid-elevation form of living primarily on the mountain slopes below 1,000m a.s.l. and reaching highest densities at about 300m [2]. Pangolins are found mainly in agricultural fields [3]. They once commonly existed throughout the lowland Taiwan in the late nineteenth century and mid twentieth century [4–6], but commercial harvesting for the domestic markets of traditional medicine and game meat, and for international trade in leather in 1950-1970 had caused their population collapsed island-wide to a very low level hence harvesting probably was no longer profitable. It was estimated that as many as 60,000 individuals were harvested annually during this period [6].

Commercial harvesting and exportation was ban by the government totally in the mid-’70s [3,7] and the relief from the high hunting pressure likely had improved the pangolin’s status in Taiwan later on [7,8]. However, although this measurement eradicated the harvest for international trade, but the demand for domestic markets apparently still existed. An estimation of more than 2,000 individuals were sold in game restaurants annually in mid-‘80s and the price for a live pangolin could be as high as USD 300 [8].

It was not until the Taiwanese government promulgated the new Wildlife Conservation Act in 1989 that the domestic consumptions were significantly suppressed. As a result, a slow and steady increase in pangolin number was observed in many locations in the recent years [9–11], unlike in other pangolin’s extant countries where local extirpation, mainly caused by intensive poaching for illegal trade, were common [12–15].

From 1993 to 2009, a total of 117 Formosan pangolins were rescued by the Endemic Species Research Institute (ESRI), central Taiwan, among which 82.9% of them were unhealthy upon arrival and required serious medical care. These records unveiled that 50% of the morbid pangolins that they rescued were injured due to gin (or foothold) traps and 23% were malnutrition, while animal attack and collision also caused serious trauma in some cases [10]. Gin traps are commonly used by farmers in Taiwan for pest control and arbitrary hunting for small mammals. Due to the high occurrence of damaging non-target animals, including invaluable wildlife, gin trap was banned from manufacturing and selling by Animal Protection Act of 2011.

Additionally, pathological findings of dead pangolins were reported by [16] from northern Taiwan and they found pneumonia and gastric ulcers to be the primary cause of mortality. Recently, Khatri-Chhetri et al. [17], from southeastern Taiwan, also found high prevalence of Hepatic and respiratory lesions in the dead bodies they examined and proposed this might be a result of long-term exposure to toxic environment. As until today pesticides and herbicides are also commonly used in the farmlands in Taiwan.

In general, although poaching is not an important issue to the Formosan pangolin, but it seems like that there are other anthropogenic threats impeding their recovery. Herein we presented two sets of information to consolidate the leading causes of mortality and morbidity of this species in the wild. One set of information came from the long-term radio tracking project in Taitung, southeastern Taiwan, during 2009-2017. The second set of information came from the pangolin rescue data procured by the Pingtung Rescue Center for Endangered Wild Animals (PTRC) during 2006-2017.

## Methodology

### Ethics statement

Ethics approval was granted by the laboratory animal center, National Pingtung University of Science and Technology (NPUST). Pangolins were captured under the licence with permission granted by the Taiwan Forestry Bureau (permit numbers 0980129850, 0991616024, 1011701139, and 1031700176) as required by the Wildlife Conservation Act, 2013. Anaesthesia and blood sampling were done with guidelines (see [18], for detailed procedures).

### Radio-tracking

Radio-tracking was conducted in the southern part of the isolated Coastal Mountain Range, Taitung County, southeastern Taiwan (22° 90’N, 121°18’ E). It is one of the areas in Taiwan where a stable pangolin population can be found. Pangolin density here is estimated to be 12.8 ind./100ha [19]. The study area is approximately 1,000 ha in size with elevation range between 100m and 700m a.s.l. The climate is generally tropical weather with hot wet season (April-November) and somewhat cooler and dryer winters (December-March). The landscape of the study area is highly fragmented, including secondary forest, bamboo forest, grassland, managed plantation, and agriculture land inlaid each other.

During 2009-2017, a total of 47 pangolins were radio-tagged with the permission issued by the Taiwan Forestry Bureau (Permit numbers: 0980129850, 0991616024, 1011701139, 1031700176 and 1050143346). Three models of radio transmitters were used, they were Telonics MOD-125 (53g with active mode; 932 E. Impala Ave. Mesa, Arizona, USA) and ATS R2030 and ATS R2020 (24g and 12g with active mode respectively; ATS, Inc. PO Box 398 470 First Ave. N. Isanti, Minnesota 55040). Transmitter selection was dependent on the body weight of the pangolin tagged where it was less than 1.5% of the body weight in all cases. Radio transmitters were attached on a scale of tail near hip. After releasing back into the wild, animals were checked for their activity signals frequently, sometime every few days, using telemetry receiver (TR4; Telonics) and a directional H-antenna (RA-2AK or RA-23K) to confirm their locations and status.

All individuals were assigned to one of the three age groups according to their body weight. Those less than 1.5kg were classified as juveniles for both sexes, while 3kg and 4kg were used to be the cutting line between sub-adults and adults for female and male respectively.

Among the 47 radio-tracked individuals, 20 were adults and 27 were sub-adults when tagged; two of them missing the original gender information, the rest were 25 females and 20 males. Un-expected incidents occurred for 22 of them during the study, of which 9 lost their signal and never been recovered, and the rest 13 individuals, involved 14 cases, were found dead or in danger by the research team. These cases provided the insight of causes of Formosan pangolin morbidity and mortality in the wild.

### Rescue data

During 2006-2017, PTRC, in southern Taiwan (22°39’N 120°36’E), rescued a total of 131 (58 females, 73 males) pangolins majority from the southern Taiwan (83.6%), especially from the southeastern Taiwan (43.9%). Their medical records included age, gender, weight on arrival, month of arrival and cause if injured.

Diagnoses were classified into morbidity categories that included trauma (encompassing limb injuries, tail injuries and collisions), other medical conditions (such as abscessation, diarrhoea and emaciation) and unknown or undetermined. The medical diagnosis was performed by veterinarians. All cases received complete physical examination. Diagnosis of the diseases or anomaly was based on signs such as evidence of infective agents (inflammatory cell aggregates, suppuration and mucopurulent discharges), and diagnostic tests such as complete blood count, serum chemistry, plasma total protein to classify emaciation and complete body X-ray to categorize trauma. Tentative clinical diagnoses were classified as undetermined or unknown.

Furthermore, many individuals were categorized healthy when arrived PTRC and were excluded from the above categories in the study. Healthy pangolins were incautiously caught by people and sent to PTRC probably due to increased awareness among the public about this cryptic mammal and popularity achieved by the rescue centers that encourages public to bring any individual they come across to the rescue centers for further assistances.

## Results

### Radio-tracking

Among the 14 cases (Table 1), number of females showed no difference between the two age groups (5:4), while much more sub-adult males were found in predicament than adult males (4:1). Case 12 and 13 belongs to the same individual. Except the first 2 cases without complete basic information, almost all of them (10 cases, 83.3%) emanated in the dryer season and the other 2 cases happened very close to the dryer season (April and October respectively); none occurred during the core wet season. The average tracking length for the known cases was 217.3 days (N=12, SD= 246.3; range: 45~869 days), majority of them happened much less than one year after being tagged (Table 1).

**Table 1.** Detailed description of bizarre mortality in fourteen radio tracked individuals of the *Manis pentadactyla pentadactyla*.

| Case No | Sex | Trap and tag |  | Died or in danger |  | Duration of tracking (days) | Description |
| --- | --- | --- | --- | --- | --- | --- | --- |
|  |  | Weight (kg) | Age group | Date | Weight (kg) |  |  |
| 1 | F | —* | Adult | 2009/—/— | > 4.00 | < 1 year | It was found stuck in hollow tree trunk and died. |
| 2 | F | — | Sub-adult | 2009/—/— | < 3.00 | < 1 year | It was found stuck in hollow tree trunk and died. |
| 3 | F | 1.65 | Sub-adult | 2010/12/15 | — | 45 | It was found died on the ground surface with unknown cause |
| 4 | M | 2.20 | Sub-adult | 2011/04/17 | 2.50 | 66 | It was found killed by gin trap and its body was handed over to the researcher. |
| 5 | F | 1.90 | Sub-adult | 2012/12/24 | — | 40 | Its tag was found detached by sharp device, pangolin was disappeared. |
| 6 | M | 2.15 | Sub-adult | 2012/12/25 | 3.80 | 468 | It was live-captured when crossing the road and handed over to researcher. |
| 7 | F | 2.10 | Sub-adult | 2013/01/15 | 2.10 | 69 | It was found died in resting burrow with unknown cause. |
| 8 | M | 3.65 | Sub-adult | 2013/02/13 | 3.00 | 98 | It was found trapped in gin trap, and was rescued and survived. |
| 9 | M | 2.60 | Sub-adult | 2014/03/07 | — | 51 | Its tag was found detached by sharp device, pangolin was disappeared. |
| 10 | F | 3.90 | Adult | 2014/03/10 | 3.60 | 367 | It was found died in resting burrow with collapsing. |
| 11 | M | 2.15 | Sub-adult | 2015/02/05 | 2.10 | 142 | It was found died in resting burrow with collapsing. |
| 12 | F | 1.85 | Sub-adult | 2015/02/20 | 4.00 | 869 | It was found stuck in a foraging burrow and was rescued. (LF28) |
| 13 | F | 4.00 | Adult | 2015/10/10 | 4.00 | 232 | It was found missing left hind limb in picture taken by camera trapper, which most likely caused by gin trap, and survived according to picture taken afterward. (LF28) |
| 14 | F | 2.80 | Sub-adult | 2016/02/09 | — | 160 | Its tag was found detached by sharp device, pangolin was disappeared. |

\* missing data.

Except Case 3 and 7 (Table 1), where causes of death could not be determined, accident happened while moving or digging into the tree or ground burrows occurred the most in our records (5 cases, 41.7%). Such fatal encounters were discernible in all ages (3 adults, 2 sub adults). Only one of the pangolin (Case 12) was still alive when found, suggested a high mortality can occur when they are stuck in the burrow. Three cases (Case 4, 8, 13) were found damaged by gin trap, and 3 other cases (Case 5, 9, 14) ascertained that the radio tag was detached from the pangolin by non-natural force, which were most likely being removed (poached) by local people when they run into these pangolins. Local people do have the opportunity to catch live pangolin by accident (e.g. Case 6). Of course, pangolins could also be caught by gin trap first and then removed (poached) by local people without being reported.

### Rescue data

Our record showed that the rescue was somehow male-biased (F:M=44.3:55.7), and involved mainly sub-adults and adults (Table 2). Number of pangolins arrived varied significantly in different months (X^2^= 25.72, df=1, p<0.005), with highest in May-June, followed by July-August and November-December with both sexes showed the similar seasonal pattern (Fig. 1). Among the 131 rescued individuals, 57 (43.5%; 31 Females, 26 males) were healthy required only proper care to recover from stress and they all were being released within 1-2 weeks from where they were discovered.

**Fig. 1.**
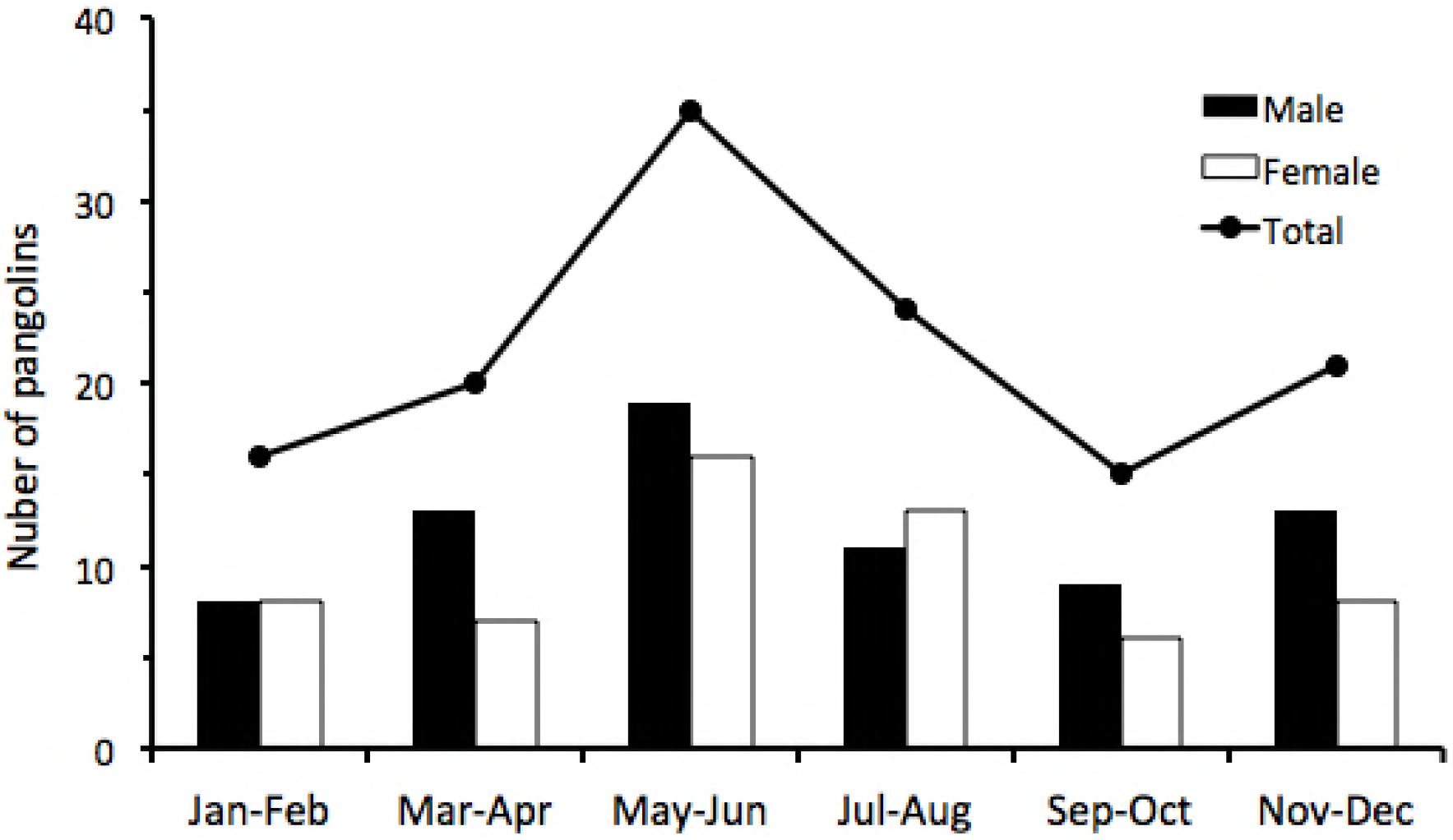
Bi-monthly arrival of Formosan Pangolin to PTRC during 2006 to 2017. Number of pangolins arrived in different months, with highest in May-June with both sexes showed the similar seasonal pattern.

**Table 2.** Number of male and female rescued by PTRC according to the age groups.

| Age groups | Female |  | Male |  | Sum |  |
| --- | --- | --- | --- | --- | --- | --- |
|  | n | % | N | % | N | % |
| Juvenile | 8 | 13.8 | 4 | 5.5 | 12 | 9.2 |
| Sub-adult | 29 | 50.0 | 39 | 53.4 | 68 | 51.9 |
| Adult | 21 | 36.2 | 30 | 41.1 | 51 | 38.9 |
| Total | 58 | 100.0 | 73 | 100.0 | 131 | 100.0 |

Compared with the sex ratio of the total rescued individuals, there were significantly more males (63.5%) in the morbid group (X^2^= 0.182, df= 1, p>0.1). The age-group structure, however, showed no difference between total and morbid pangolins for both female (X^2^= 0.613, df= 2, p>0.1) and male (X^2^= 0.778, df= 2, p>0.1), suggested that the morbidity is not age-depended. Trauma (73.0%) was the most prominent reason of morbidity (Table 3). Cases of trauma involved limb injuries (77.8%) were all caused by gin traps, while those with tail injuries (20.4%) were most likely caused by dog bite; 1 individual experienced collisions caused by vehicles. The rest were other medical conditions that include emaciation, diarrhea and abscessation (Table 3). There was no gender difference in the morbid types distribution (X^2^= 0.44, df= 3, p>0.1).

**Table 3.** Causes of morbid in Formosan pangolin rescued by the PTRC.

| Morbid | Female | Male | Sum |
| --- | --- | --- | --- |
| Trauma | 20 | 34 | 54 |
| *Limb injury* | 14 | 28 | 42 |
| *Tail injury* | 6 | 5 | 11 |
| *Collisions* | 0 | 1 | 1 |
| Other medical conditions | 4 | 10 | 14 |
| Injury data missing | 3 | 3 | 6 |
| Total | 27 | 47 | 74 |

Majority of the rescued pangolin were released back into the wild, but 1/10 of them died eventually (Table 4). Releasing sites were safe environments or protected area close to where they were found. Again, there was no gender difference in the result of the rescue action (X^2^= 0.52, df= 2, p>0.1).

**Table 4:** Success rate of rescued pangolin cases.

|  | Female |  | Male |  | Sum |  |
| --- | --- | --- | --- | --- | --- | --- |
|  | n | % | n | % | n | % |
| Pangolins released into wild | 53 | 91.4 | 63 | 86.3 | 116 | 88.5 |
| Pangolins for long term captivity | 0 | 0.0 | 1 | 1.4 | 1 | 0.8 |
| Pangolins died | 5 | 8.6 | 9 | 12.3 | 14 | 10.7 |
| Total | 58 |  | 73 |  | 131 |  |

## Discussion

From the two sets of information presented here, it clearly appears that the damage caused by the gin trap is the prime threat to the life of the Formosan pangolin, it comprised more than half (56.8%) of the morbid cases in the rescue records (Table 3) and 25.0% in the radio-tracking cases (Table 1). The previous data from the ESRI in central Taiwan also unveiled similar pattern that 61.8% of morbid pangolins rescued by them were induced by gin traps [10]. As mentioned earlier that the production and selling of gin traps had been legally banned in Taiwan since 2011, however Lin [20] did a survey recently in rural Taiwan and found 23.1% (= 9/39) of the hardware stores they interviewed still offered gin traps to the farmers. Unregulated installation of gin traps in agriculture lands, as a commonly used pest control measurement, is also a major life threatening factor to the endangered leopard cats (*Prionailurus bengalensis*) in rural Taiwan [21,22]. Although strengthening the law enforcement is urgently required, provide farmers with animal friendly techniques to substitute the present inhuman pest control devices should also be mandatory.

Our assessment done directly by radio-tracking in the wild brought in staggering results, which was obscure did not show in the rescue records, that pangolin has high accidental mortality during its daily activity. Pangolins being fossorial in nature, they are proficient diggers and being named as *Chuan-Shan-Jia* (means Mountain Excavation Armor) in Chinese. However, the surprisingly high risk during excavation for food searching or resting burrow maintenance have subjected the pangolins to unexpected mortality caused by being stuck in tree hollows or underground burrows. Such incidences were prominent during the dryer season (Table 1), when digging for termites became inevitable because supply of ants, their primary source of food, was scares [23–27]. The paucity of food during the dryer season inflicted loss of body weight [28], hence probably also poorer bodily condition, that might further reduce their capability to escape from such an unprecedented danger. Lin [20] reported that the average length of feeding burrow was 120.1±69.5cm, while that for the resting burrow can even reach 201.6±94.8cm. Different landform or land use practices might affect the properties of soil, and some might increase the possibility of being trapped, therefore, this kind of fatal accidents can cause significant damage to the local population [29]. Furthermore, the fragmented geography in southeastern Taiwan is mainly dominated by various agricultural practices that supersede the natural composition of the landform hence increasing the risk of non-natural deaths in Formosan pangolin.

Although the number of dog attacks was comparatively less, but its impact cannot be overlooked. *Canis familaris* have outnumbered all the canids collectively and has perturbed various wildlife communities through the act of predation [30,31]. Dogs are also responsible for posing serious threat other than dog bites. Lenth et al. [32] observed the impertinent intrusion of dogs in exploring new territories led to the restriction in the motility of various wildlife, as well as have depicted temporal shift in their pattern of activity. Therefore, dogs might also indirectly affect the motility pattern of pangolin making more discernible to other predators like humans. For example, the change of route due to presence of free ranging dogs might dispose pangolins to other injuries from the traps since they may enter the modified landforms supplemented with trapping devices.

Moreover, dogs are believed to be affected by the ticks especially belonging from Ixodidae family. The same family of ticks is known to affect the pangolins as well [33,34]. Ticks especially in their nymph stage are known to affect the medium sized and large mammals and can be a serious threat to the health of the individual since they carry various pathogens [35]. In search of potential host to complete the life cycle ticks perform questing that could affect pangolins since they position themselves under the scale causing infections in them [33,35] because pangolins have very sensitive immune system [36].

The other medical conditions that incorporates diarrhea, emaciation and abscessation contributes to another reason for mortality and morbidity in Formosan pangolin. Autopsy records disseminated that gastric ulcers are very prominent in this species [16,18] and that might act as a contributing factor for diarrhea in them. Furthermore, pangolins might face the challenge of unavailability of food resources due to extermination of land resources by unwanted human activities. The dwindling of sufficient food resources subjects Formosan pangolin towards emaciation as it perturbs the fundamental dietary requirements generating stress which is correlated with the generation gastric ulcers.

Susceptible to injuries especially during the summer months from May to August (Fig. 1), as sub-adults emerge from the weaning period and maternal care are expected to be prepared to find new unoccupied territory. The similar pattern had also been found with the previous reported rescue records [10]. In the course of finding new regions, pangolins might come across the areas subjected to anthropogenic disturbances steering them towards more injuries due to their explorative behavior [37]. A higher rescue incidence also occurred during the winter months from November to January (Fig. 1), which is the reproduction season of the Chinese pangolin [38]. As such, more traveling, hence more accidents are expected.

The amount of injuries experienced by males are higher than females in both the records from ESRI and PTRC might be due to much larger home range in males [20], so they were more active than the females leading males to distend themselves to certain areas [16]. The similar behaviors of males have been observed in other species and males are subjected to be brought into rescue centers more often [39,40]. However, it is also might due to females are more fragile accounts for their higher mortality rate to any kind of injuries, because they are smaller and lighter than males, therefore less female will be found surviving from injuries in the wild. The same reason might also explain why there were only a few juveniles in the rescue records.

Despite the appalling situation generated by trapping devices and agricultural practices, the joint efforts of different rescue centers along extensive sharing network of knowledge across Taiwan about this cryptic mammal turned the daunting situation into pangolin amicable. This communal effort led to enhanced understanding and awareness among the diaspora culminating towards the declining rate of mortality of Formosan pangolin. The rescue centers ESRI and PTRC are the epitome of this alliance of teamwork where they could bring down the rate of mortality from 35.9% to 10.7% (Table 4). Furthermore, tremendous success records were observed since rate of releasing pangolin into wild also increased from 60.7% to 88.5%.

The two records depict contrasting observation as well, the ESRI projected the percentage of rescued healthy individuals were less than the healthy pangolins brought to PTRC (17.1% vs 43.5%). The increment in the percentage might have occurred due to increased awareness among the public and popularity gained by the rescue centers that might encourage people to turn in the pangolin more than before. It also might be due to the expansion of pangolin’s population, so more individuals come in contact with human concentrated areas during locomotion. The data obtained states that rescue centers have acted as a prophylactic measure for recovery of Formosan pangolin from ancillary injuries as well as has acted as an institution promulgating public education, and research that have directly contributed to in-situ conservation.

For conservation of endangered species it is fundamental to understand given species mortality factors. Through this paper we were able to track down some of natural and unnatural reasons for the loss of the Formosan pangolin and some of them newly discovered. However, despite of there are so many factors threatening their life, the Formosan pangolin’s population, unlike in all other pangolin range countries, is stabilized and even started to increase in many areas in the past decades [9,10]. This provided a strong evidence that poaching for illegal markets, which is no longer a threat to the Formosan pangolin, alone can cause the extirpation or endangered status for a species as in other range countries.

## Acknowledgments

We would like to express our gratitude for the field assistants Man-Rong Yu, Xiang-Hua Hu, Jia-Hui Lin and Ya-Wen Leu for their tireless and unfathomed support in this extensive field work. We appreciate and thanks Taitung Forest District Office, Taitung County Government, Taitung County Police Bureau (LuanShan station) and Formosan Pangolin Conservation Association for their logistic supports. We are also grateful to the staff of the Pingtung Rescue Center for Endangered Wildlife for investing time and taking complete care of rescued pangolins and meticulously taking a record of them. Also thanks Taitung Forest District Office (Grant number: 104-737-1), Ministry of Science and Technology (Grant number: NSC 101-2621-M-020-006, NSC 102-2621-M-020-004, NSC 102-2621-M-110- 004) and The Mohamed Bin Zayed Species Conservation Fund (Grant number: 13256024) providing monetary assistance.

## Data Availability Statement

All relevant data are within the paper.

## Author contributions

**Conceptualization:** Nick Ching-Min Sun; Kurtis Jai-Chyi, Pei

**Data curation:** Wen-Chi Lin, Meng-Jou Ch, Chen-Chih Chen

**Formal analysis:** Nick Ching-Min Sun; Bharti Arora

**Funding acquisition:** Kurtis Jai-Chyi, Pei; Nick Ching-Min Sun

**Investigation:** Nick Ching-Min Sun; Bharti Arora; Jing-Shiun Lin

**Methodology:** Kurtis Jai-Chyi, Pei; Jing-Shiun Lin

**Supervision:** Kurtis Jai-Chyi, Pei

**Writing-original draft**: Nick Ching-Min Sun

**Writing-review & editing:** Kurtis Jai-Chyi, Pei; Bharti Arora

